# Orienting to Polarized Light at Night—Matching Lunar Skylight to Performance in a Nocturnal Beetle

**DOI:** 10.1101/366583

**Authors:** James J. Foster, John D. Kirwan, Basil el Jundi, Jochen Smolka, Lana Khaldy, Emily Baird, Marcus J. Byrne, Dan-Eric Nilsson, Sönke Johnsen, Marie Dacke

## Abstract

For polarized light to inform behaviour, the typical range of degrees of polarization observable in the animal’s natural environment must be above the threshold for detection and interpretation. Here we present the first investigation of the degree of linear polarization threshold for orientation behaviour in a nocturnal species, with specific reference to the range of degrees of polarization measured in the night sky. An effect of lunar phase on the degree of polarization of skylight was found, with smaller illuminated fractions of the moon’s surface corresponding to lower degrees of polarization in the night sky. We found that South African dung beetle *Escarabaeus satyrus* (Boheman, 1860) can orient to polarized light for a range of degrees of polarization similar to that observed in diurnal insects, reaching a lower threshold between 0.04 and 0.32, possibly as low as 0.11. For degrees of polarization lower than 0.23, as measured on a crescent moon night, orientation performance was considerably weaker than that observed for completely linearly-polarized stimuli, but was nonetheless stronger than in the absence of polarized light.

**Summary Statement:** A degree-of-polarization threshold for orientation behaviour is reported for nocturnal dung beetle *Escarabaeus satyrus* in the context of measurements showing changes in the degree of polarization of skylight with lunar phase.

## Introduction

Many animals use the sun or the moon to orient their movements (Papi & Pardi, 1963; Frisch, 1967; Schmidt-Koenig, 1990; Dacke *et al*., 2014). When the sun is not directly visible, its position can be estimated from the pattern of polarized skylight produced by Rayleigh scattering (Strutt, 1871) in the upper atmosphere. The use of these solar skylight polarization cues during daytime has been studied in numerous species (reviewed in: Wehner, 2001; Zeil *et al*., 2014), and similar behaviour has been shown to extend into twilight (Dacke *et al*., 1999; Dacke *et al*., 2003a; Freas *et al*., 2017). Orientation to polarized lunar skylight has, to date, only been demonstrated in two species of night-active dung beetle found in southern Africa, *Scarabaeus zambesianus* and *Escarabaeus satyrus* (Dacke *et al*., 2003b; Dacke *et al*., 2004, Dacke *et al*., 2011). These beetles sculpt a dung ball, which they roll away from the dung pat, to a distance where it can then be buried and consumed without interference from other beetles. In order to travel in a straight line, these species use celestial cues as compass references (Dacke *et al*., 2011). In contrast to the closely related diurnal dung beetle *Kheper lamarcki, E. satyrus* orients to polarized light cues in preference to the observable position of the moon (el Jundi *et al*., 2015a) and may be specialised to detect the faint lunar skylight that emanates from a crescent moon (Smolka *et al*., 2016).

### Degree of Polarization of Lunar Skylight

Linearly-polarized light, such as solar- and lunar skylight, is defined by two properties: its angle and degree of polarization. Both properties vary across the sky as a function of angular distance from the sun or moon, forming a pattern that indicates its position. When either sun or moon are at the horizon, this pattern is an ambiguous cue indicating either an eastward or westward celestial body. When the sun or moon is above or below the horizon, and the observer is aware of whether it is before or after sunset/moonset, this pattern can be interpreted unambiguously. The angle of polarization is the axis along which the greatest proportion of a light beam’s electric field strength oscillates, while the degree of (linear) polarization is that beam’s intensity in this axis as a proportion of the beam’s total intensity. As a result, degree of polarization can be considered a measure of signal strength for an animal using the skylight pattern of angles of polarization as an orientation cue. Relative to the sun or moon’s position, this angle-of-polarization pattern is similar across a range of conditions (Gál, *et al*., 2001; Hegedüs *et al*., 2007; Barta *et al*., 2014; Wang *et al*., 2016). In contrast, the degree of polarization decreases as a result of cloud cover and atmospheric turbidity (Labhart, 1999; Hegedüs *et al*., 2007; Wang *et al*., 2016) as well as being affected by light pollution on moonlit nights in urban areas (Kyba *et al*., 2011). Since lunar skylight is far dimmer than solar skylight, contributions to celestial light from other sources can reduce the maximum observable degree of polarization in the night sky, when these light sources have either a low degree of polarization (*e.g.* zodiacal light, integrated starlight, skyglow: Kyba *et al*., 2011) or a malaligned angle-of-polarization pattern (*e.g.* solar skylight: Barta *et al*., 2014). For *E. satyrus*, which relies heavily on skylight cues, a failure to detect such weakly-polarized skylight could mean failure to orient, which in the worst case would result in returning to the high-competition region around the dung pile.

It has been suggested that the field cricket *Gryllus campestris* has adapted a low threshold (degree of linear polarization = 0.05) that permits the detection of the weakly polarized skylight often observed in its natural environment (Labhart, 1996, 1999). An assessment of the daytime skylight cues available near the crickets’ collection site, using an artificial polarization-sensitive “neuron”, suggested that degrees of polarization are typically quite low, as a result of haze and cloud cover, with median values of 0.13 and 0.23 in the solar and antisolar halves of the sky respectively (Labhart, 1999); as compared with values in excess of 0.60 measured in clear skies (Horváth *et al*., 2014). While the low intensity of lunar skylight presents a challenge for the detection of polarized skylight at night, the superposition eyes of *E. satyrus* are well adapted to detect lunar skylight cues, and they orient well even under very dim conditions (Dacke *et al*., 2011; Smolka *et al*., 2016). Nevertheless, if the combination of moonlight with other, unpolarized, celestial light results in weakly-polarized skylight, then *E. satyrus* would require a degree-of-polarization threshold that is low enough to match the typical range found in its geographic distribution.

### Detection Thresholds for Polarization

The thresholds for detection of polarized skylight previously estimated for insects have varied both between species and experimental paradigms. In a series of studies involving the field cricket *G. campestris* (Labhart, 1996; Henze & Labhart, 2007), elliptically-polarized light was used to investigate the degree of linear polarization (DoLP) threshold. The threshold for polarization-opponent interneurones in the optic lobes was estimated at DoLP = 0.05 (Labhart, 1996). In behavioural experiments, reorientations of tethered walking crickets viewing a rotating stimulus with a degree of linear polarization as low as 0.03 were consistently greater than those measured for a circularly-polarized stimulus, though not significantly different after controlling for multiple testing (Henze & Labhart, 2007). Interestingly, neuronal responses to rotating elliptically-polarized light in the central brain of desert locust *Schistocerca gregaria* indicated a threshold as high as DoLP = 0.30 (Pfeiffer *et al*., 2011). The behavioural threshold reported for honeybees (*Apis mellifera*) is intermediate between the two examples given above. Bees were still observed to orient their “waggle dances” well to spots of skylight with degrees of polarization as low as 0.10 (Frisch, 1967, p403–404), while dances for degrees of polarization >0.07 were described as “not completely disoriented”.

In this study we investigated the polarization of lunar skylight across different lunar phases at field sites in South Africa near *E. satyrus*’ typical range. We also tested the robustness of *E. satyrus*’ orientation behaviour to decreases in the degree of linear polarization, allowing us to compare the beetle’s behavioural threshold with the measured properties of lunar skylight.

## Methods

### Polarization Imaging of Skylight

Lunar skylight is many orders of magnitude dimmer than its solar equivalent, making it more challenging to measure via photopolarimetry. The first published study of the polarization of lunar skylight used a series of images recorded through different polarizer orientations onto photographic film (Gál *et al*., 2001), since commercially available charge-coupled device (CCD) sensors were deemed insufficiently sensitive at the time. Just a decade later, a dark-current-corrected CCD-based system was successfully employed to compare lunar skylight between urban and rural areas (Kyba *et al*., 2011), also following a serial-imaging protocol. Recently, interactions between solar- and lunar-skylight polarization were observed by comparing the untransformed signal from three separate CCD cameras (Barta *et al*., 2014), each measuring a different angle of polarization, avoiding time-series artefacts. In this study we used a single camera with a complementary metal–oxide–semiconductor (CMOS) sensor, which was dark-corrected and calibrated to compensate for lens distortion and nonlinearities in CMOS chip sensitivity prior to estimation of polarization across the sky.

In order to assess the typical range of polarization states in lunar skylight, ‘polarization images’ of the sky were created for lunar phases ranging from full moon to new moon. Photographs were recorded at the game farm ‘Stonehenge’ (near Vryburg: 26°23’56”S, 24°19’36”E) and at Thornwood Lodge (near Bela-Bela: 24°46’08”S, 28°00’52”E), both in South Africa, at the Neusiedler See Biological Station (47°46’05”N 16°46’03”E) in Austria, and the Finnish Meteorological Institute Arctic Research Centre (67°21’60”N 26°37’42”E) near Sodankylä, in Finland, using a digital camera (D810: Nikon Corp., Japan), fitted with a fisheye lens (Sigma 8 mm F3.5 EX DG: Sigma Corp., Japan) and a filter holder (CA483-72: Sigma) with a polarizing filter (WR 72mm: Sigma). The system was previously calibrated to allow us to obtain an estimate of the absolute spectral radiance values for the red, green and blue channels (Nilsson and Smolka, in prep.). Moon fullness data for each night was retrieved from the U.S. Naval Observatory’s website (http://aa.usno.navy.mil/). To create a single polarization image, 25 photographs were recorded: One set of five photographs with the camera aimed directly at the zenith, and then one set each with the camera aimed at the horizon to the north, east, south and west. Between each image, the polarizer was rotated anti-clockwise (relative to the camera’s viewing direction) by 45°, thus recording a set with the transmission axis at 0°, 45°, 90°, 135° and 180° to its starting orientation (west–east when the camera faced the zenith and to the camera’s right when it faced the horizon).

From the raw images, we calculated an estimate of absolute spectral radiance in the blue range of the visual spectrum. Images were then filtered to simulate a 4° half-width Gaussian filter to approximate the upper limit of visual resolution in *E. satyrus* (Nilsson and Smolka, in prep; Foster *et al*., 2017). Polarization parameters were estimated for each direction by comparing the images taken at different filter orientations. An estimate of unpolarized intensity, in the absence of measurement error, may be calculated as the sum of the 0° and 90° images; or of the 45° and 135° images; or of the 90° and 180° images (*N.B.* since the 0° image preceded the 90° image, while the 180° image was taken afterwards, we used both to obtain a more accurate estimate of average unpolarized intensity). The differences in radiance between the 0° and 90° images and the 45° and 135° images were taken as Stokes parameters S_1_ and S_2_ respectively, and the average of the sums of each pair was used to estimate total intensity (Stokes parameter S_0_). From these values the angle and degree of linear polarization were calculated for each pixel. Finally, we combined the images obtained for the different directions into a full hemispheric image by assigning each pixel the value of the directional image whose visual axis was closest, avoiding large off-axis viewing angles through the polarizer, and the associated intensity artefacts (Foster *et al*., 2018), where possible.

### Orientation to Polarized Light

Beetle collection and the behavioural experiments were carried out at “Stonehenge”, 70 km North-West of Vryburg, North-West Province, South Africa. Beetles were kept in sand-filled boxes, where they were fed with cow dung *ad libitum*. Prior to each experiment, beetles were removed from their boxes and allowed to sculpt balls of cow dung and roll them with an unobstructed view of the moonlit sky. Experiments were conducted between the 11th and 15th of November, 2016, under clear conditions during which the fraction of the moon’s illuminated surface visible ranged between 86% and 100%.

### Polarized light stimulus

Stimulus light was provided by four UV-emitting 18W fluorescent bulbs (LT-T8 Blacklight Blue: NARVA Lichtquellen GmbH, Germany) in addition to eight ‘cool white’ LEDs (DDW-UJ2- TU1-1: Roithner LaserTechnik GmbH, Austria), because visible-spectrum light was found to be necessary to maintain beetle activity. The light source was directed through a stack of seven diffusing filters (Fig. 1 A), each constructed from white shading cloth (Euro-Serre shade: Willab AB, Sweden) attached to a 6 mm-thick plate of UV-transmissive acrylic (Perspex, U.K.). For all conditions this stack also contained a polarizer (HNP’B: Polaroid Corp., U.S.A.). For the “unpolarized” and “maximally-polarized” conditions this stack was reconfigured to include a sheet of translucent drafting paper that acted as an additional diffuser (Fig. 1 B), but somewhat reduced the subsequent UV content of the stimulus. These diffusers allowed the reduction of the degree of linear polarization of the stimulus light without the introduction of elliptical polarization (as is the case when using a circular polarizer), which in some rare cases can be converted back to linearly polarized light in the animal’s eye (Shurcliff, 1955; Choiu *et al*., 2008; Templin *et al*., 2017). In this study we can therefore disregard elliptical polarization, and its potential conversion to linear polarization, and instead refer to degree of linear polarization (DoLP) as degree of polarization wherever it was manipulated or measured. A similar stimulus was used in a recent study to investigate the polarization threshold of aquatic springtail *Podura aquatica* (Egri *et al*., 2016).

**Figure 1.**
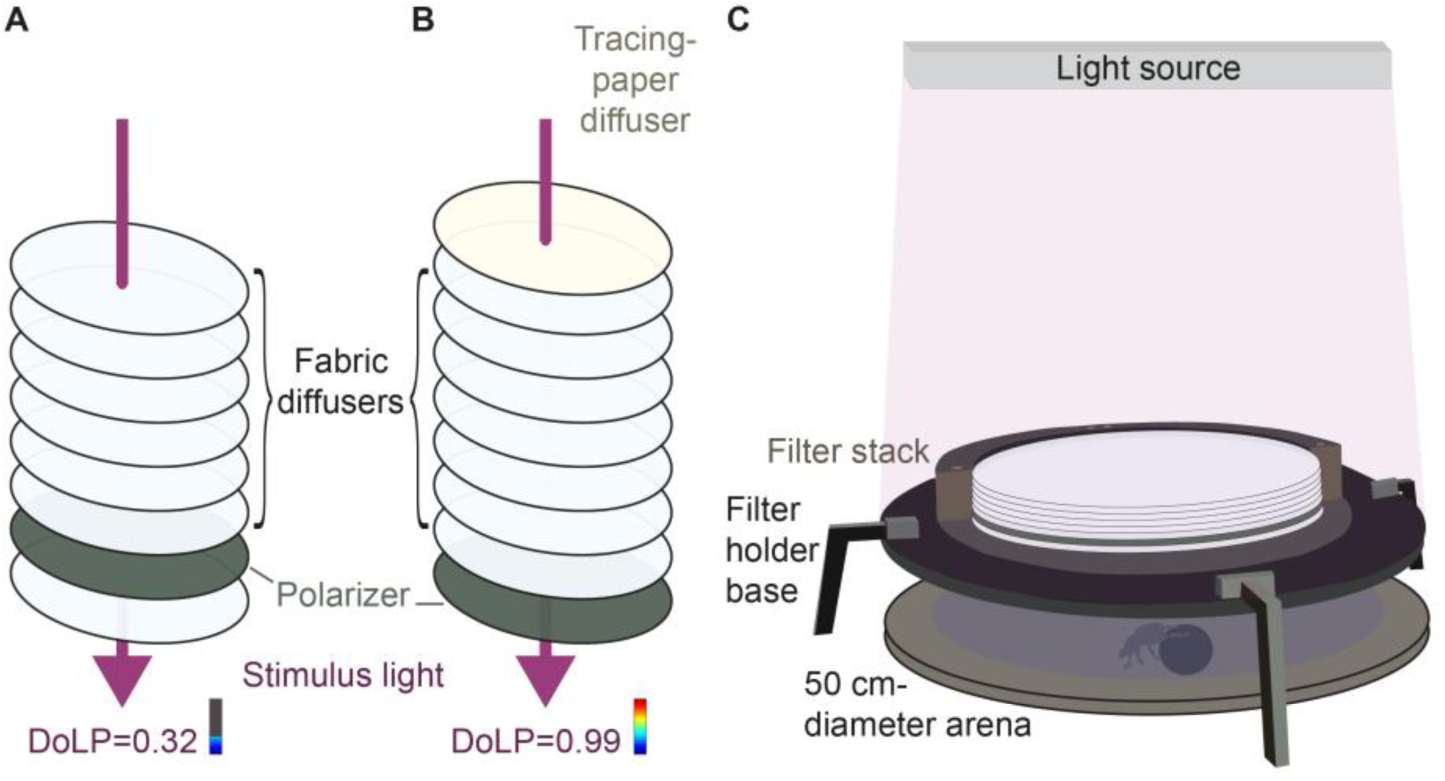
The filter stack used in orientation experiments. The filter stack is shown arranged so as to produce (A) the polarized stimuli and (B) the “maximally-polarized” and “unpolarized” controls. (C) shows the arrangement of the filter stack during experiments. (A) Unpolarized light from a fluorescent lamp passed through seven acrylic-mounted fabric diffusers and a polarizer, so that the number of diffusers before and after the polarizer in the light path determined the degree of polarization of stimulus light. In the arrangement shown (6 diffusers before the polarizer, 1 diffuser after the polarizer) stimulus light would have a degree of polarization of 0.32, while if the whole filter stack were inverted stimulus light would have a degree of polarization of <0.02 (1 diffuser before the polarizer, 6 diffusers after the polarizer). (B) The addition of a drafting-paper diffuser ensured that degrees of polarization of 0.99 and ≈0 could be produced by one filter stack, depending on whether it was upright (“maximally-polarized”: as shown) or inverted (“unpolarized”). (C) The filter stack was suspended 12 cm above a 50 cm diameter arena from which the beetle viewed stimulus light transmitted through it, but not the light from the fluorescent lamp itself, which was shielded from view by the filter holder’s base (outer casing of the filter stack shown in cross-section to reveal the filters inside).

The intensity of each condition (Supplement S1.1) was measured from the position of the arena’s centre using a calibrated irradiance probe (cosine corrector: CC-3-UV-T; light guide: P600-2-UV-VIS; spectrometer: QE65000; all produced by Ocean Optics Inc., Dunedin, USA). Polarization was measured at the same position using a UV-transmissive calcite linear polarizer (Glan-Thompson; GTH5M-A: Thorlabs GmbH, Germany) coupled to a spectrometer (FLAME-S-UV-VIS) via a light guide (P1000-2-UV-VIS; Ocean Optics). Spectra were recorded for four polarizer orientations in order to estimate Stokes parameters S_1_ and S_2_ (Foster *et al*., 2018) and each measurement repeated ten times and averaged (Supplement S1.2) to minimise the effects of sensor noise (Tibbs *et al*., 2018). Prior to Stokes parameter estimation, median spectrometer response for each polarizer orientation was weighted by the absorption spectrum of an insect photopigment with a maximum absorbance at 365 nm, calculated using the Stavenga template (Stavenga, 2010), and integrated across the region of the spectrum from 380–450 nm. This was done to limit the influence of regions of the spectrum for which the spectrometer received insufficient signal and those outside of the range detectable by a dung beetle UV-sensitive photoreceptor.

To ensure that minimally and maximally oriented behaviour were observed, stimuli for which no measurably polarized light reached the animal (DoLP ≈ 0, “unpolarized”; UV irradiance 325–400 nm = 8.5 × 10^11^ photons cm^−2^ s^−1^) or in which light was strongly-polarized and of equivalent brightness (DoLP = 0.99; “maximally-polarized”; UV irradiance = 3.4 × 10^11^ photons cm^−2^ s^−1^), were produced (Fig. 1 B). Following this, stimuli with degrees of polarization of 0.32, 0.11, 0.04 at UV irradiances of 4.5, 9.0, 4.6 × 10^11^ photons cm^−2^ s^−1^, respectively, were tested. For each of these experiments, the filter stack was arranged so that it could be inverted and the order of the filters in the light path reversed. As a consequence, when the filter stack was arranged to produce a polarized stimulus, it could be rapidly inverted between trials to produce a degree of polarization of <0.02 (at UV-irradiances 9.0, 4.5, 8.9 × 10^11^ photons cm^−2^ s^−1^ respectively). This alternation occurred after every second individual, so that each individual experienced either a “polarized” or “control” condition.

### Orientation Experiments and Analysis

For each experiment, beetles were allowed to roll a dung ball to the edge of a 50 cm diameter circular arena, where its bearing was noted. Once the stimulus had been rotated by 90°, each beetle was then replaced at the centre of the arena and allowed to roll to the edge a second time. The 90° rotation of the stimulus was subtracted from the second of these two headings, to give a measure of heading relative to the stimulus’ transmission axis. Because the response of a polarization-sensitive photoreceptor to a given angle of polarization follows a 180° periodicity, accurate orientation under this scenario is expected to follow an axial-bimodal pattern, with some well-oriented individuals changing their heading by 180°. To account for this axial pattern, headings were doubled prior to further analysis (Batschelet, 1981). The length of the resultant mean vector for each pair of headings was taken as a measure of orientation precision. In this formulation a perfectly-oriented individual, orienting using polarization alone, would reorient by 0° (relative to the stimulus’ transmission axis), producing a mean vector of length 1. Following this transformation, a Prentice test (Prentice, 1979) was used to test for differences in mean vector length between each experiment’s polarized test and ‘unpolarized’ control conditions. Mann-Whitney rank sum tests were used to compare mean vector lengths between the polarized and control conditions for each condition-pair *post hoc*, with a Bonferroni-Holm correction for multiple testing.

## Results

### Skylight Polarization across Lunar Phases

We found that gibbous moon skylight (93–98% fullness; Fig. 2 A–B) measured in rural South Africa reached degrees of polarization as high as 0.6–0.7, similar to values reported for sunlit skies (Berry *et al*., 2004; Horváth *et al*., 2014) but much greater than the levels, around 0.27, for lunar skylight measured in Europe (Supplement S2.1 A; 80% illuminated). We also found that the maximum degree of polarization for other lunar phases was lower, with degree of polarization corresponding to moon fullness (Fig. 2 G). When the moon was close to the horizon, modal degrees of polarization within 60° of the zenith were 0.43 for an intermediate fullness (two days before last quarter: 80% illuminated) and 0.23 for a crescent moon (20% illuminated). Modal degrees of polarization measured during a moonless and an overcast night with a gibbous moon (67% illuminated) were around 0.06 and 0.08 respectively. Though the movements of a few stars and planets across the image set produced small regions with artificially high (false) “degrees of polarization”, measured degrees were very low across most of the sky in the absence of lunar skylight. The lack of alignment of the angles of polarization in adjacent sky regions (Supplement S2.2 E–F) suggests that non-zero values result from measurement error, rather than true sources of atmospheric polarization (*e.g.* airglow and light pollution on the clear night and transmitted lunar skylight on the overcast night).

**Figure 2.**
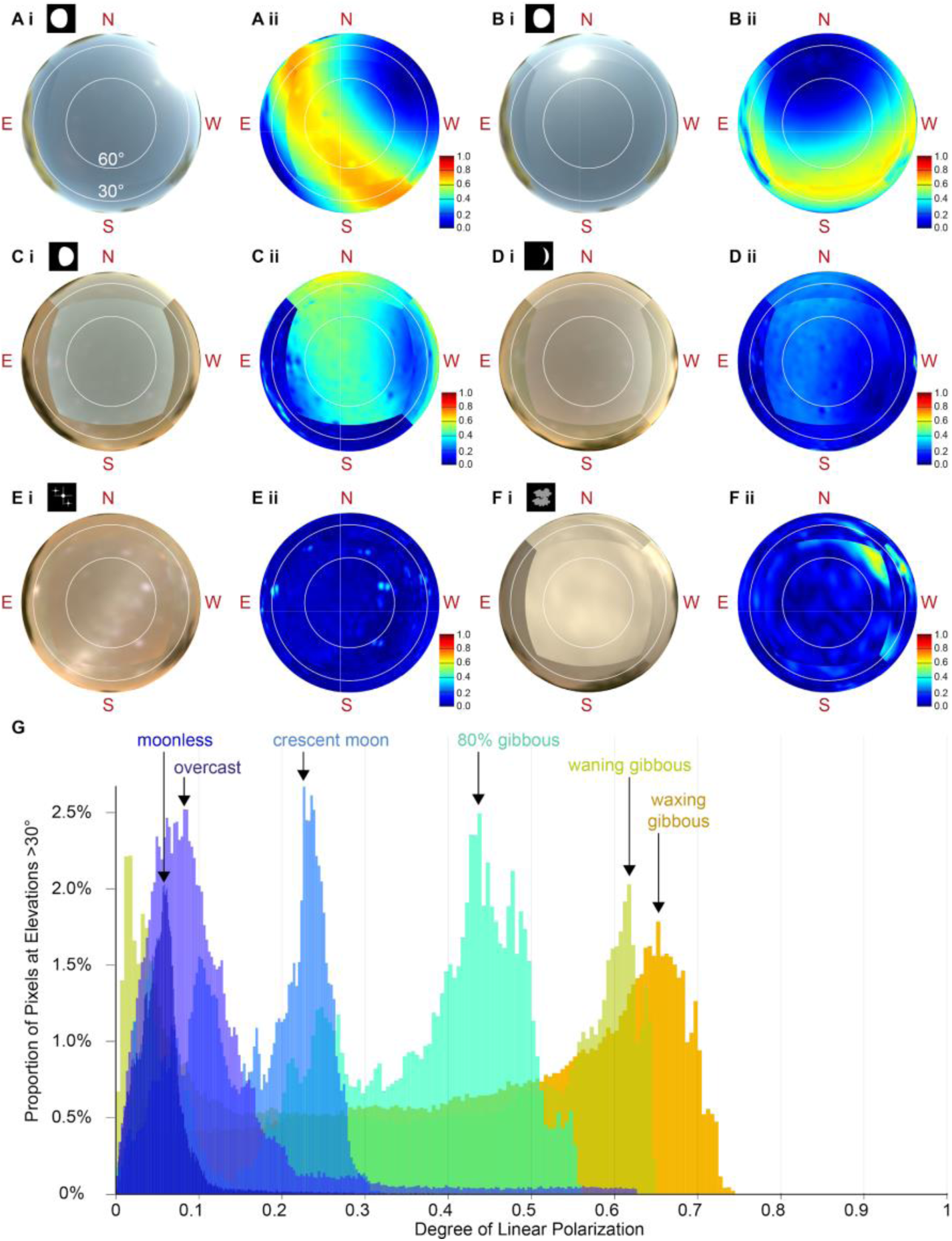
Spectral radiance (i) and degree of linear polarization (ii) images of night skies. Image sets recorded before (A) and after (B) full moon, and under gibbous moon (C), crescent moon (D), moonless (E) and overcast gibbous moon (F) conditions in rural South Africa. All images are displayed on a radial azimuth-elevation grid, with concentric white rings at elevations of 30° and 60° (A i). Azimuth values are relative to local magnetic north. Radiance images have been linearised and normalised to their brightest value (excluding the moon itself in A and B). Estimated degree of linear polarization is relative to the colour map at the bottom right of each panel; redder hues indicate high degrees of polarization, intermediate degrees of polarization are greener hues, and degrees of polarization approaching zero (i.e. unpolarized) are darker blue hues. In (A) the moon was near the north-western horizon, causing the maximum degree of polarization band to cross the sky from north-east to south west, at an angular distance of 90° from the moon itself. In (B) the moon was at 45° elevation to the north, and the band of maximum degree of polarization ringed the sky at 45° elevation in the south, crossing the horizon in the east (where it is obscured by trees below 30° elevation) and west. In (C) and (D), the moon was near the western horizon and the maximum-degree band crossed north-south and near the zenith. In (E) the moon was too far below the horizon (≥18°) to produce measurable lunar skylight. (F) shows an overcast sky lit by a gibbous moon, for which lunar skylight was not detectable through the thick cloud cover. Bright moonlight (0–45° elevation to the north-west) enhanced motion artefacts that artificially inflated degree of polarization estimates in that region. (G) shows histograms of degrees of polarization measured for each camera pixel at elevations >30° (excluding vegetation near the horizon). Between the crescent moon (light blue) and gibbous moon (orange) measurements, the modal degree of polarization (indicated by black arrows) increased as a function of moon fullness from 0.23 to 0.65. *N.B.* For the waning gibbous moon, the upper mode (0.62) is indicated, rather than the lower mode (0.02) which corresponds to the region around the moon itself. (A), (B), (C) and (E) were measured near Vryburg, and (D) and (F) were measured near Bela-Bela.

### Orientation Behaviour

For the highest degree of polarization (DoLP = 0.99) beetles were well oriented in an axial-bimodal distribution (axial mean vector length: ρ= 0.82, 0.71–0.98; mean, 1st and 3rd quartiles) consistent with previous observations for this species when presented with artificial polarized stimuli (el Jundi *et al*., 2015a). Mean vector lengths differed between the polarized and control conditions (Prentice *χ*^2^ = 19.617, d.f. = 4, *p* <0.001), and for degrees of polarization greater than 0.04 beetles were significantly better oriented (DoLP = 0.99: W = 1745, *p* = 0.001; DoLP = 0.32: W = 1041.5, *p* = 0.020) or show some indication of improved orientation (DoLP = 0.11: W = 963, *p* = 0.078) compared with the unpolarized controls.

The difference in the concentration of the axial distribution of heading changes (Fig. 3) as a function of degree of polarization was notable, with each decrease in the degree of polarization producing a corresponding decrease in axial mean vector length distributions (1st and 3rd quartiles: ρ_0.99_ = 0.71–0.98, ρ_0.32_ = 0.63–0.97, ρ_0.11_ = 0.50– 0.97, ρ_0.04_ = 0.40–0.87). To investigate this effect further, we modelled changes in the concentration parameter of a von Mises distribution, κ, as a function of degree of polarization (Fig. 4). Following initial inspection of the trend, as well as previous studies of the relationship between degree of polarization and response strength (*e.g.* Labhart, 1996; Glantz and Schroeter, 2006), we chose a complimentary log-linear relationship between κ and degree of polarization: log(κ) = *β* log_10_(DoLP). The base ten was chosen for straightforward examination of the relationship, such that a slope of 1 would indicate a tenfold increase in the magnitude of orientation precision between degrees of polarization of 0.099 and 0.99 (corresponding roughly to the lowest and highest values for which oriented behaviour was observed). A Bayesian generalised linear model was implemented in the Stan language (Carpenter *et al*., 2017), using the package brms 2.3.1 (Bürkner, 2017) in R 3.5.0 (R Core Team, 2018). Individual heading biases were accounted for as individual-level effects on mean heading, and effects of trial number, individual and test night on log(κ) were included as random effects. The fitted model had the formula, log(κ) = 3.28 + 1.41*x*, where *x* = log_10_(DoLP) − log_10_(0.02). While the model’s credible intervals were very broad distribution (Fig. 4), at degrees of polarization greater than 0.11 the model’s interquartile range (Fig. 4: red shaded area) diverged from that of its intercept (Fig. 4: blue shaded area), further suggesting that polarization may contribute somewhat to orientation performance above this value. Taken together, our sky measurements and behavioural data suggest that the degree of polarization of lunar skylight during a crescent moon may be close to the threshold for oriented behaviour.

**Figure 3.**
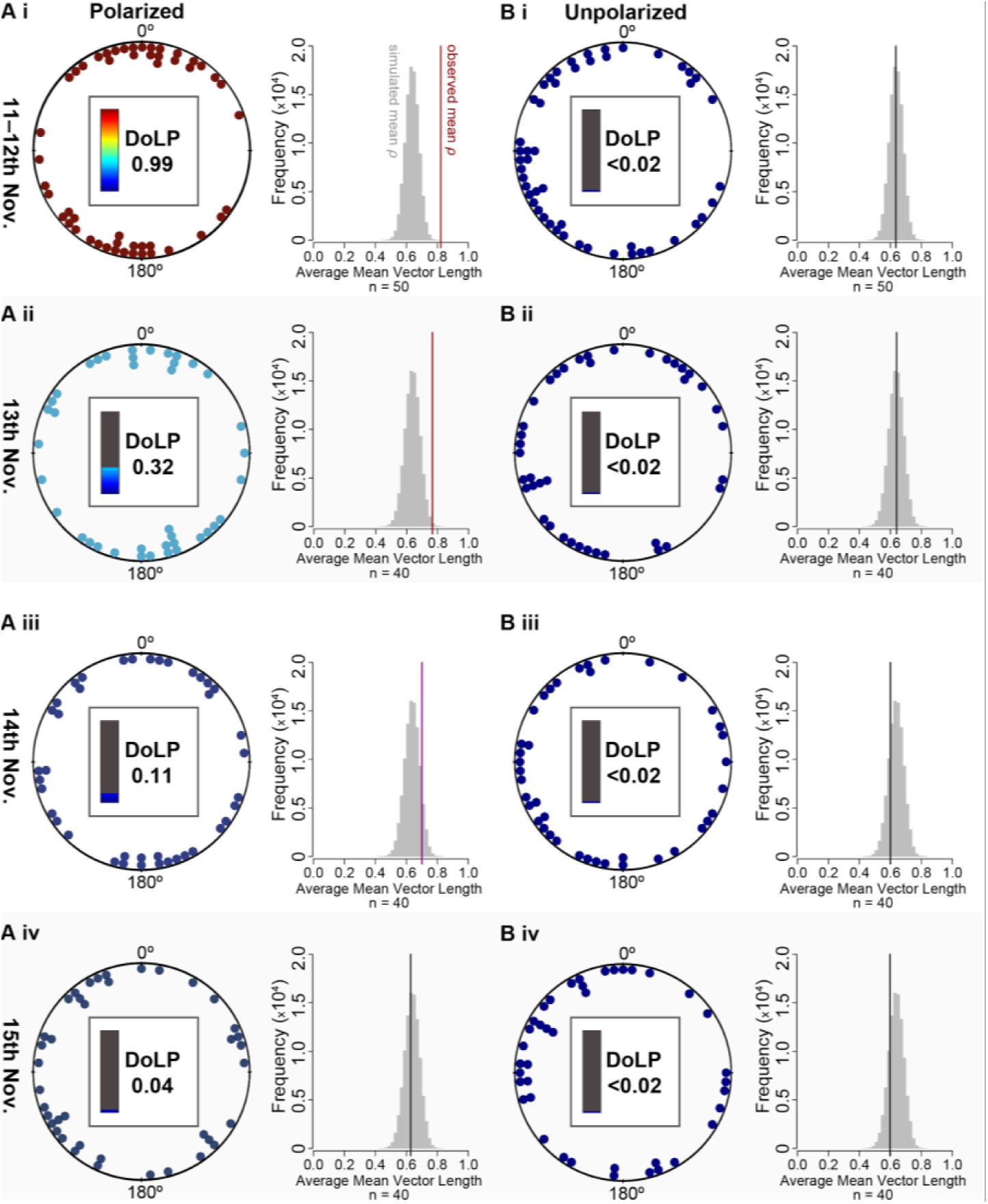
Orientation change for stimulus light with different degrees of linear polarization. Differences in heading between two sequential trials, between which the stimulus’ angle of polarization (AoP) was rotated by 90°, shown relative to stimulus angle of polarization (*i.e.* AoP_East_ – AoP_North_ – 90°). Each point represents relative orientation change for one individual, for one pair of trials, and points are colour coded to correspond to the colours used to plot degree of linear polarization for lunar skylight (see Fig. 2). Degree of linear polarization is shown in bold type at the centre of each circle. To the right of each circular plot is a simulated distribution (grey histogram) of average mean vectors for the same number of disoriented (κ = 0) individuals. Red, magenta or black lines indicate average mean vector lengths for each condition that correspond to either the top 5%, top 10% or lower 90% of 100 000 simulations, respectively. Column (A) shows the ‘polarized’ conditions, decreasing in degree of linear polarization from (A_i_) (DoLP = 0.99) to (A_iv_) (DoLP = 0.04) and column (B) shows the unpolarized condition for each experiment (DoLP ≈ 0). For degrees of polarization of 0.99 (A) and 0.32 (A_ii_), orientation precision was significantly greater than in the corresponding unpolarized control (B_i_ and B_ii_).

**Figure 4.**
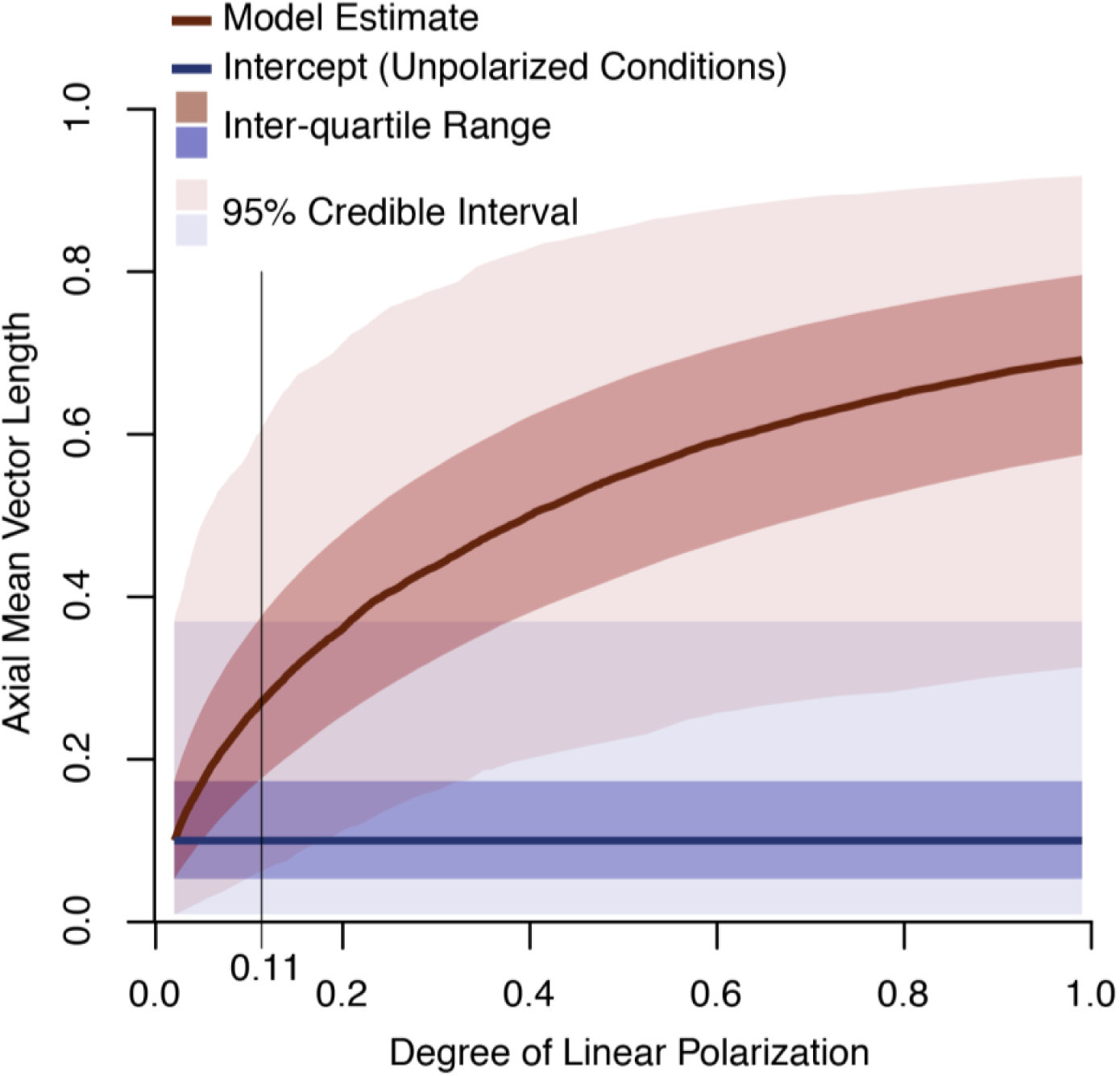
A fitted model for the relationship between degree of polarization and orientation precision. The red line indicates the fitted regression line (mean estimate) of the model for all stimuli. The pink shaded region indicates its 95% credible interval and the red shaded region indicates its inter-quartile range (50% of model estimates). The blue line shows the model’s intercept at the control condition (DoLP < 0.02) and the lighter blue and darker blue shaded areas indicate its 95% credible interval and inter-quartile ranges respectively. Model estimates of circular concentration parameter *κ* have been transformed to mean vector length ρ. The overlap of the inter-quartile ranges of the model (red shaded area) and its intercept at the control conditions (blue area) suggests that orientation performance may be aided by polarization for degrees of polarization as low as 0.11.

## Discussion

### Polarization of Lunar Skylight

On all moonlit nights the highest degree of polarization was measured in a band at approximately 90° from the moon (Fig. 2), mimicking that of the sunlit sky (Berry *et al*., 2004; Horváth *et al*., 2014), as found for previous studies of the polarization of lunar skylight (Gál *et al*., 2001; Kyba *et al*., 2011; Barta *et al*., 2014; Tang *et al*., 2016). Although the serial-imaging protocol we employed can lead to the accumulation of motion artefacts, particularly during long exposures, these effects appeared to be limited, for the most part, to a small number of bright stars (Fig. 2 E) and brightly-lit cloud edges (Fig. 2 F) in the absence of lunar skylight. Our results indicate both that lunar skylight can be nearly as polarized as solar skylight (DoLP ≥ 0.6) and that its degree of polarization is modulated by lunar phase. One previous study of lunar skylight polarization also reported a lower maximum degree of polarization for a gibbous moon than a full moon (72% illuminated: DoLP ≈ 0.15; 78% illuminated: DoLP ≈ 0.13; 100% illuminated: DoLP ≈ 0.25; Barta *et al*., 2014). Since the intensity of lunar skylight corresponds to the fraction of the moon’s illuminated surface observable (approximated as spectral irradiance = 1 – [cos(illuminated fraction × π/2)]^0.29^: Palmer & Johnsen, 2015; Kieffer & Stone, 2005), we propose that any reduction in the intensity of polarized lunar skylight relative to the intensity of other light sources usually decreases the degree of polarization of celestial light, which is a composite of the two.

In general, the degrees of polarization in South African night skies reported here are greater than those reported in previous studies conducted in central Europe (0.36 ± *σ* 0.02, Elstal, Germany: Kyba *et al*., 2011; 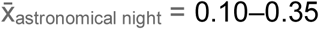, Szombathely, Hungary: Barta *et al*, 2014). The two sites in South Africa at which we measured lunar skylight were at higher altitudes (Vryburg: 1202 m, Bela-Bela: 1137 m), have a semi-arid climate and low light pollution, all of which may play a role in permitting strongly-polarized lunar skylight to reach a terrestrial observer unhindered and undiluted. Using the same method in central Europe produced similar estimates of degree of polarization to those found in previous studies (mode = 0.27, Illmitz, Austria: Supplement S2.1 A).

Clear-sky polarization patterns can be predicted as a function of atmospheric turbidity and wavelength (*e.g.* Wang *et al*, 2016). When atmospheric turbidity is low—for example, as a result of low concentrations of water droplets—the degree of polarization of skylight is greater, but is lower in the UV wavelengths than in the blue, the former of which *E. satyrus* are thought to use to detect polarized light (el Jundi *et al*., 2015a, 2016). Since our analysis focussed on the camera’s ‘blue’ channel (full-width at half maximum of sensitivity: 420–505 nm) it is possible that we do indeed overestimate the degree of polarization available to a dung beetle. The UV content of a moonlit sky may be lower than the equivalent proportion for sunlight (−14% photons_350–400 nm_:photons_400–450 nm_, calculated from: Johnsen et al., 2006), while other sources of celestial light, such as airglow (Benn & Ellison, 1998) and starlight (Johnsen *et al*., 2006) have more similar intensities in the blue and UV regions. Future studies could use a combination of polarimetric spectroscopy and photopolarimetry to more accurately determine how wavelength, turbidity and lunar phase interact to produce different degrees of polarization of lunar skylight, and how this might impact nocturnal and crepuscular insects that detect polarization in either the UV or blue regions of the spectrum. This may be of particular importance given the recent dramatic increases in the intensity of skyglow from anthropogenic light pollution (Falchi *et al*., 2016) and its potential to reduce the degree of polarization of lunar skylight (Kyba *et al*., 2011) below the detection thresholds for dung beetles and other nocturnal and crepuscular arthropods that may orient using polarized lunar skylight.

### Polarization Sensitivity Thresholds

For this study, we expanded the analysis methods used for neuronal recordings in crickets (Labhart, 1996) and locusts (Pfeiffer *et al*., 2011) to fit response curves to circular concentration of (re-)orientation behaviour. We report an estimated threshold for orientation behaviour in *E. satyrus* at between degrees of polarization of 0.04–0.32, modelled at 0.11, under dim-light conditions, which is broadly comparable to those found for other insects in daylight (0.05 for *G. campestris*: Labhart, 1996; Henze & Labhart, 2007; 0.10 for *A. mellifera*: Frisch, 1967; 0.30 for *S. gregaria*: Pfeiffer *et al*., 2011). *E. satyrus* orients to skylight both during the full moon, when skylight remains bright throughout the night, and when only a thin crescent of the moon’s surface is illuminated, providing dim lunar skylight for a brief period directly after dusk or before dawn (Smolka *et al*., 2016). The compass neurons in *E. satyrus*’ central complex have been shown to weight the predominant angle of polarization (a proxy for the skylight polarization pattern) more strongly than the position of a light spot (such as the sun or moon), in contrast to diurnal dung beetle *K. lamarcki*, which weights the light spot’s position more strongly (el Jundi *et al*., 2015a). *K. lamaracki* is thought to combine information from multiple skylight cues, including intensity gradients (el Jundi *et al*., 2014) and spectral gradients (el Jundi *et al*., 2015b, 2016), and *E. satyrus* may also rely on a combination of skylight cues when the moon becomes obscured. It is possible that, to achieve the impressive orientation precision observed on nights lit by a crescent moon (Smolka *et al*., 2016), *E. satyrus* builds on the performance observed here for polarization cues in isolation by integrating information from multiple skylight cues.

### Threshold Analysis

In our behavioural experiments, we observed an increase in orientation accuracy as a function of degree of linear polarization, from which we derive our estimate of *E. satyrus*’ polarization threshold. We note, however, that methods for defining a polarization threshold have varied somewhat across the literature. In *G. campestris*, the threshold for electrophysiological recordings was defined as the point at which firing-rate modulation was above the 95% confidence interval for responses to circularly-polarized light (DoLP = 0) and darkness (Labhart, 1996), whereas for behavioural experiments walking direction modulations were compared non-parametrically with the circularly-polarized control. Firing-rate modulation was also calculated for *S. gregaria*, but the mean vector length derived from these values was instead chosen for comparison with a circularly-polarized control, again identifying a threshold at the condition for which all recordings were outside of the (Gaussian) 95% confidence interval (Pfeiffer *et al*., 2011). In each case, these definitions avoided implying orientation capacity for conditions where responses to polarized and unpolarized light were at all similar, but did not take trends across the dataset as a whole into account (although regression models were reported: Labhart, 1996; Pfeiffer *et al*., 2011). In this study we attempt to use this trend to inform our estimate for the point where the responses to polarized and unpolarized light diverge. This definition allows us to extrapolate from our data, proposing a minimum degree of polarization at which any facilitation of orientation could occur, which is a form of ‘threshold’. Nevertheless, definitions based on confidence intervals, either explicitly or through statistical null-hypothesis testing (including the Prentice test and Mann-Whitney rank sum tests reported here), are liable to change with increasing sample size. A particular hindrance for this dataset was that, since only two trials were carried out per individuals, uncertainty in orientation precision at the individual level was high, resulting in broad credible intervals for the fitted model (Fig. 4; pink shaded area).

Future work might benefit from both a greater number of trials per individual and the fitting of a psychometric curve for responses to polarized stimuli (*e.g.* Temple *et al*., 2015). This type of model asymptotically approaches both the baseline rate of response, when there is no stimulus, and the maximum rate of response, beyond which increases in stimulus strength produce only infinitesimal increases in response strength. The inflection point of such a curve may be taken as an estimate of threshold (Knoblauch & Maloney, 2012; Houpt & Bittner, 2018). This has the advantage that it does not compound uncertainty in the positions of both the baseline of the response curve and its slope. Such methods have not yet been developed for circular data, although analogous measures have been used in a few recent studies (Foster *et al*., 2017; Kirwan, 2018).

### Detection Limits

While behavioural thresholds can indicate what conditions would allow animals to use polarized light in nature, they do not necessarily represent a true detection threshold in the sense of the visual system’s physiological limits. Animals may continue to detect polarized light, but discard the information, or weight it more weakly compared to other cues. Inhibition of responses to weakly-polarized light may be adaptive for skylight-orienting insects. For a polarization compass to aid in identifying the sun’s true position, the angle-of-polarization pattern of the sky as a whole must be integrated and combined with information about time of day. This would require an internal representation of the angle-of-polarization pattern that changes over the day, as has been demonstrated for the tunings of neurons in the central brain of locusts (Pfeiffer & Homberg, 2007; Bech *et al*., 2014). Evidence from both locusts (Bech *et al*., 2014) and honeybees (Rossel & Wehner, 1984) indicates that this representation best matches solar skylight in the high degree-of-polarization band 90° from the sun, where angles of polarization in adjacent sky regions are most aligned. It has therefore been suggested that weakly-polarized regions could be excluded from the sun-compasses of some species (Rossel & Wehner, 1984; Pfeiffer *et al*., 2011), since they may correspond to the regions that are poorer matches to this simplified internal representation of the angle-of-polarization pattern.

Bernard & Wehner (1977) proposed that the sensitivity (S) of a photoreceptor to a beam of light illuminating it should be proportional to,

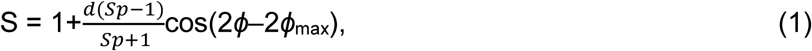

where *d* is the degree of linear polarization of the beam, *ϕ*_max_ is the angle of polarization (*ϕ*) to which the photoreceptor is maximally sensitive, and polarization sensitivity, *S*_*p*_ is the response to *ϕ*_max_ divided by the response to *ϕ*_max_±90° if *d* = 1. When the response (resulting from S) is sufficiently distinct from the response to unpolarized light, it should be possible for an eye containing photoreceptors sensitive to different angles of polarization to detect and interpret polarized light. To meet this requirement: i.) *S*_*p*_ must be greater than one, ii.) *d* must be sufficiently large, iii.) the beam must be bright enough for modulation as a function of *ϕ, d* and *S*_*p*_ to be distinguishable from sources of noise. In general, the influence of the angle of polarization (*ϕ*) might be reasonably discounted for hymenopterans and ball-rolling dung beetles, which often perform complete body axis rotations when commencing orientation behaviour. In this study we focussed on requirement (ii): that degree of polarization must be sufficiently large, taking orientation to completely linearly-polarized light as indicative that requirements (i.) and (iii.) were met.

The blue-sensitive dorsal rim photoreceptors of *G. campestris* have mean polarization sensitivity of *S*_*p*_ = 8.3 (Labhart *et al*., 1984), which would suggest that at threshold there is only a 2–4% difference (*d* = 0.03–0.05) between sensitivity to unpolarized and partially-polarized light. For the dorsal rim UV-receptors of *A. mellifera*, mean polarization sensitivity is *S*_*p*_ ≈ 5 (Menzel & Snyder, 1974), indicating a difference around threshold of 5–7% (*d* = 0.07–0.10). By contrast, a similar modal value of *S*_*p*_ ≈ 8.5 for the dorsal rim blue receptors of *S. gregaria* indicates that this difference is 24% at threshold (*d* = 0.30). Somewhat larger values of *S*_*p*_ = 15.1 and *S*_*p*_ = 7.7–12.9 were reported for dorsal rim UV receptors of diurnal and crepuscular dung beetles *Pachysoma striatum* (Dacke *et al*., 2002) and *S. zambesianus* (Dacke *et al*., 2004), respectively, and assuming similar values for *E. satyrus* would give a difference of 8–9% (*d* = 0.11) around threshold. Considering the relatively small increases in sensitivity required to elicit oriented behaviour, it is plausible that the performance of the visual systems of *G. campestris, A. mellifera* and *E. satyrus* are noise-limited at threshold, while that of *S. gregaria* appears to be inhibited by some other process. Nevertheless, since no photoreceptor recordings are currently available for *E. satyrus*, we cannot exclude the possibility that these beetles may detect but disregard degrees of polarization that fall outside the range found in the moonlit skies observable in their natural habitat.

Nocturnal and diurnal species might also face different constraints in the detection of polarized skylight. Many nocturnal species (such as *E. satyrus*) or species active during both day and night (such as *G. campestris*) pool visual signals from adjacent regions (Warrant, 1999) to increase signal-to-noise ratios through larger absolute photon catches. Such spatial pooling could lead to the combination of signals from regions with misaligned angles of polarization, and of high degree of polarization and low degree of polarization regions, reducing the maximum observable degrees of polarization while enabling more robust detection of dim skylight polarization patterns. The stimuli used in this study were 2–3 orders of magnitude dimmer than those used in most previous studies (Labhart, 1996; Pfeiffer *et al*., 2011; el Jundi *et al*., 2014), although they remain 1–3 orders of magnitude brighter than the UV component of a moonlit sky (Johnsen *et al*., 2006: Supplement S1.1). Light intensities used in this study are most similar to those for the ‘stimulus size’ experiment performed with *G. campestris* (Henze & Labhart, 2007), in which an opaque annulus surrounded a 1° diameter completely linearly-polarized stimulus (to which the crickets successfully oriented). *G. campestris* is active during both day and night, and it is possible that their impressive polarization sensitivity also allows them to orient to polarized moonlight as *E. satyrus* does. In this initial study to investigate the effects of degree of polarization on orientation accuracy in dung beetles, we have not addressed the third condition outlined above: that detection of polarization can only occur when polarized light is sufficiently bright. To more accurately define the limits for a beetle’s skylight compass, future studies should compare orientation performance across the full light-intensity and degree-of-polarization range of lunar skylight. Such information could help to predict how anthropogenic pollution can affect nocturnal arthropods, and aid in the development of solutions to mitigate them.

## Conclusions

The nocturnal dung beetle *Escarabaeus satyrus* (Boheman, 1860) has previously been demonstrated to orient to polarized lunar skylight throughout the lunar month, and appears well adapted to detect dim lunar skylight. In darkroom experiments employing dim polarized stimuli with a range of degrees of polarization, we find that *E. satyrus* remains oriented across a range similar to that reported for diurnal insects, reaching a threshold between 0.04 and 0.32, possibly as low as 0.11. We also provide measurements of the variation in degree of polarization of lunar skylight across different lunar phases, recorded in *E. satyrus*’ natural habitat.

## Supporting information

## Author Contributions

JJF, BeJ, EB and MD conceptualised the study and designed the behavioural experiments. JJF, BeJ, LK, EB, MJB and MD carried out the behavioural experiments. JJF and JS devised the polarization-imaging protocol, and JS and DE-N conceptualised and designed the sky-imaging system. JS designed the polarization-analysis software. JDK and SJ conceptualised the statistical analysis and JDK designed and carried out the statistical modelling. JJF drafted the manuscript and all authors revised the manuscript.

## Acknowledgements

The authors thank Ted and Winnie Harvey of Stonehenge game farm, and Riitta Aikio and Markku Ahponen at the FMI Sodankylä -Pallas Arctic Research Centre for their help in the field, and Therése Reber in particular for braving high towers and long Arctic winter nights to obtain measurements. We would also like to thank Camilla Sharkey for advice regarding beetle spectral sensitivity, Bob Fitak for advice on analysis and David O’Carroll for providing the drafting-paper diffuser. We dedicate this paper to the memory of Ted Harvey, whose help, patience and ingenuity have proven vital to the past decade of dung beetle orientation research, and who will be sorely missed.

## Supplement

### S1 Stimulus Calibration

**S1.1.**
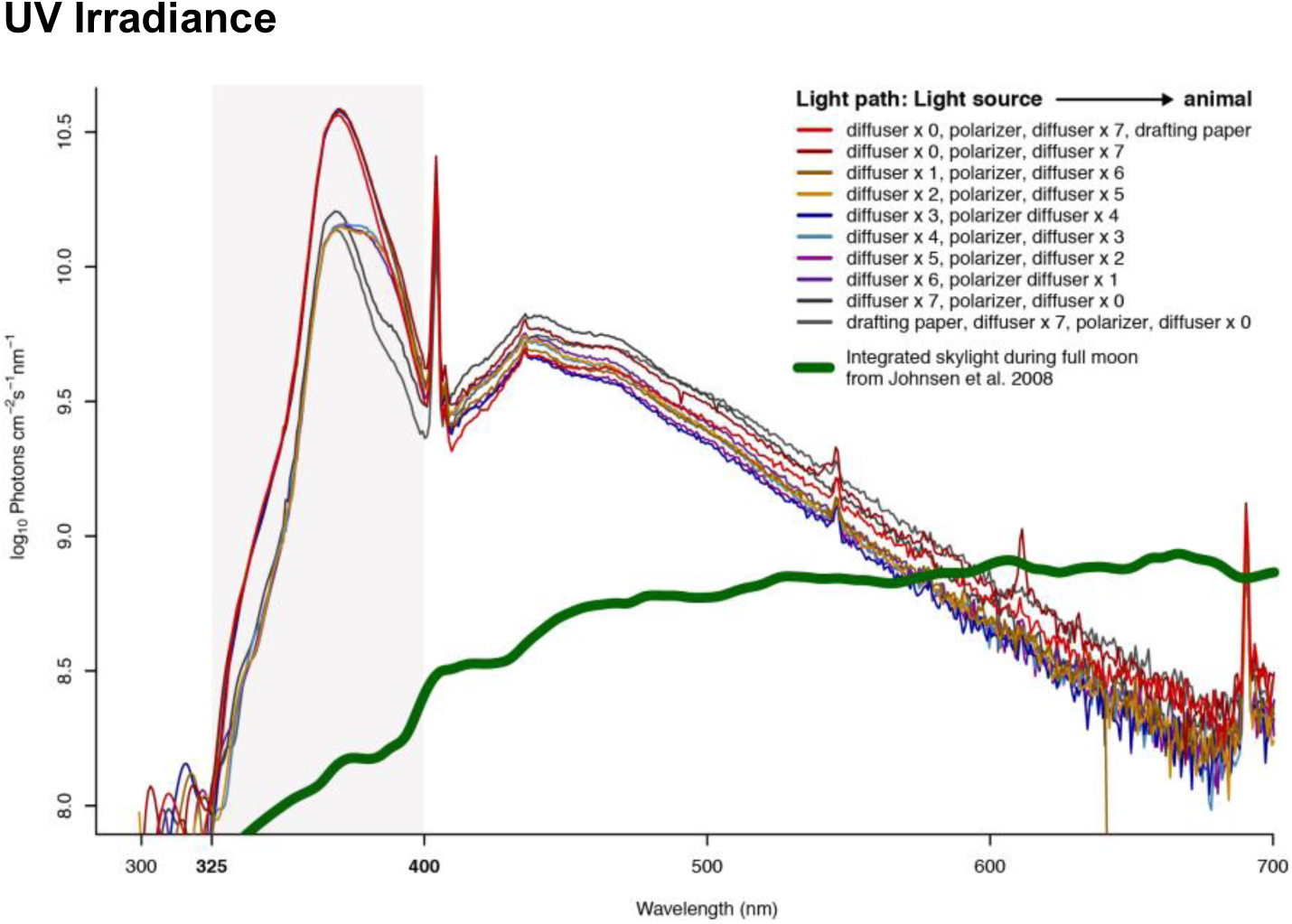
Spectral irradiances measured for each condition. Spectral irradiance was measured at the position of the arena’s centre (where beetles were placed at the beginning of each trial) and summed across the spectrum between 325 nm and 400 nm. We estimate that spectrometer signal was adequate for accurate measurements across this range and that the photoreceptors responsible for detection of polarized light in *E. satyrus* are likely to reach peak sensitivity within this region. Irradiances for each stimulus reported in the main text were determined by integrating across the spectrum between 325 nm and 400 nm (region shaded in the figure above).

**S1.2.**
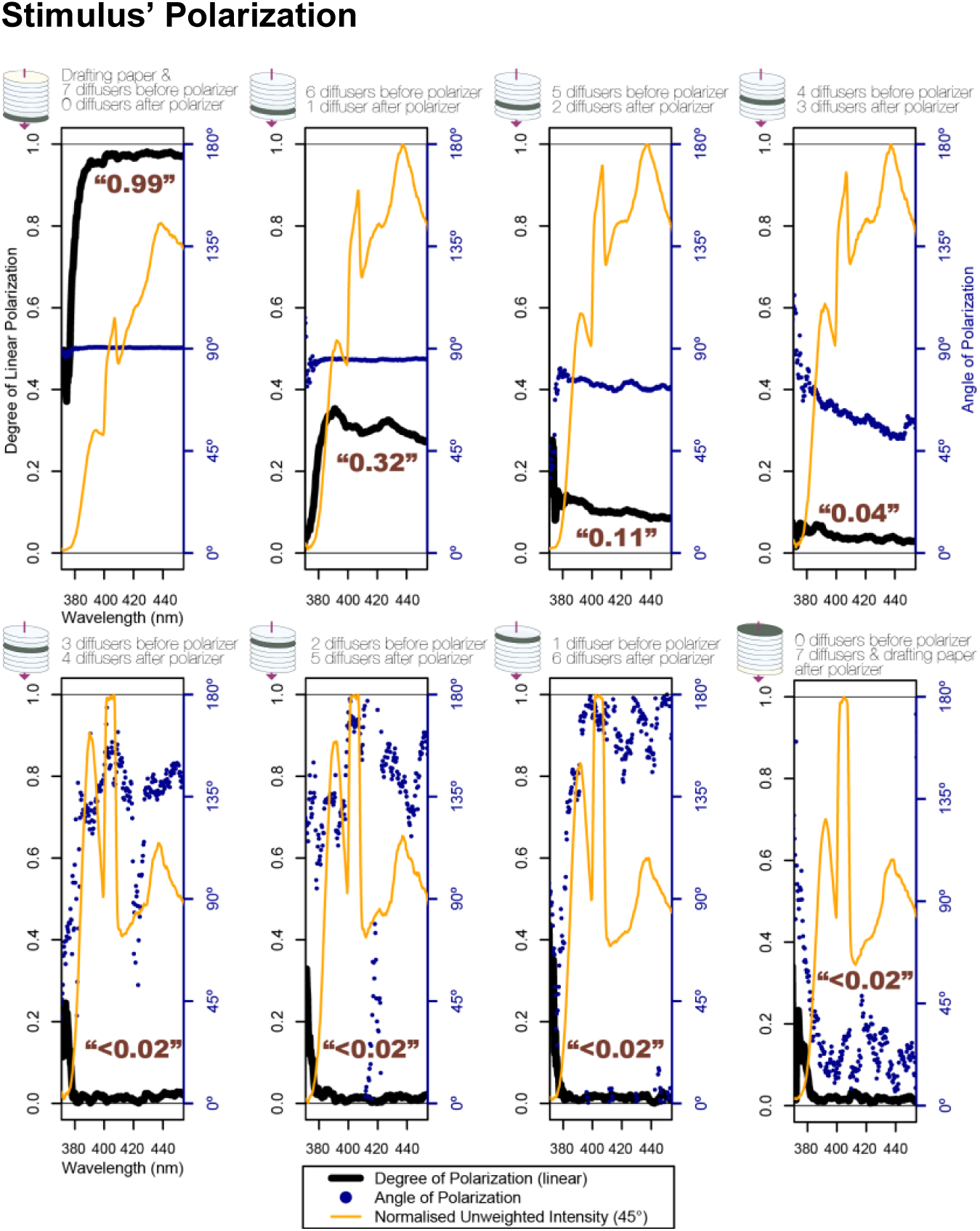
Spectral characterisation of polarization for each experimental condition. The polarization of each stimulus condition was characterised across the UV-visible spectrum using a linear polarization filter within a lens tube (to restrict off-axis light) coupled to a spectrometer. Wavelengths shorter than 380 nm not shown, since the spectrometer’s signal was extremely weak below this value. The spectrometer’s signal was weighted by the absorbance spectrum of an insect UV-sensitive opsin with a maximum absorbance at 365 nm, approximately midway between the values reported for other scarabaeids (Lord *et al*., 2016) and within the range indicated for *Pachysoma striatum* (Dacke *et al*., 2002) and *S. zambesianus* (Dacke *et al*., 2004). These spectra were then integrated across the range 380–450 nm, for which both spectrometer signal and UV-opsin sensitivity were both relatively high, to provide estimates of relative intensity for each polarizer orientation, which could then be used to calculate Stokes parameters S_0_, S_1_ and S_2_, and subsequently the degree of linear polarization.

### S2 Lunar Skylight Measurements

**S2.1.**
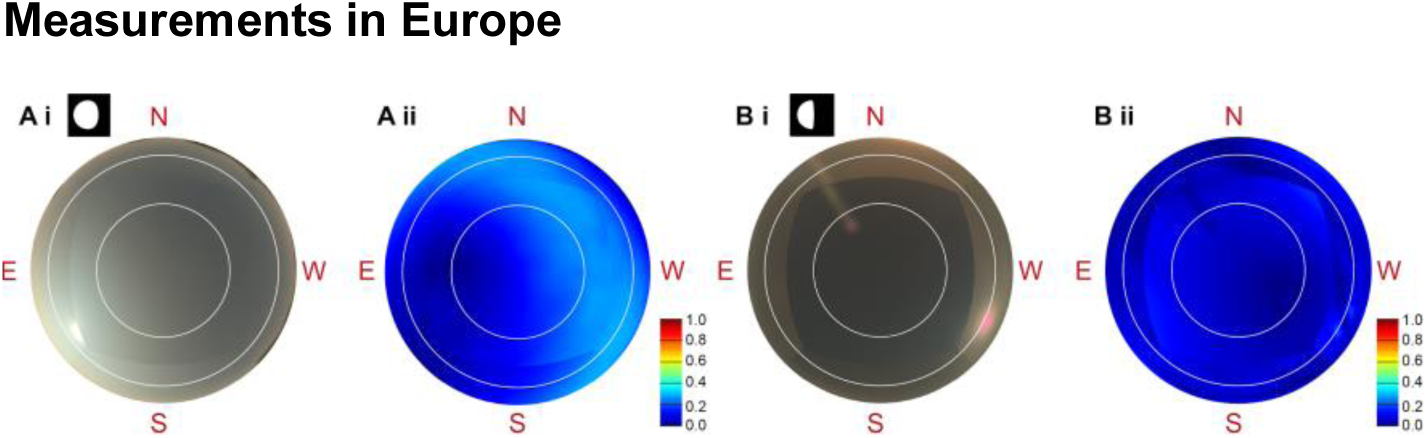
Intensity and degree of polarization images recorded for rural moonlit skies in Europe. At the Neusiedler See Biological Station (47°46’05”N 16°46’03”E) light pollution was low (A; moon: 30° elevation, south-east, 80% illuminated), and the degree of polarization in the strongly-polarized band 90° from the moon was 0.27 (modal value), similar to that reported in previous studies conducted in central Europe (Gál *et al*., 2001a; Kyba *et al*., 2011; Barta *et al*., 2014). This site has a warm, temperate-oceanic climate and is at an altitude of 117 m above sea level. Skies above the Södankylä -Pallas Arctic Research Centre in Finnish Lapland were more light-polluted (B; moon: 30° elevation, south-west, 38% illuminated), and the degree of polarization in the high-degree band was considerably lower, between 0.08 and 0.17 (mode in this band: 0.11). This value is similar to that measured in the night sky in Berlin (0.17±*σ* 0.02: Freie Universität measurement tower: Kyba *et al*., 2011), but lower than those measured at other rural sites in this study. The tower from which these measurements were recorded includes an antenna that partly blocks the view of the sky to the north-east up to 70° elevation, resulting in an unpolarized region. This tower was previously used for measurement of solar skylight during 25 h of daylight in an Arctic summer (Gál *et al*., 2001b), when maximum degrees of polarization were found to fall in the range 0.5– 0.7, as for clear skies in other regions. The site has a wet subarctic climate and is at an altitude of 179 m above sea level (the tower is 16 m above ground level, removing trees and buildings from the field of view shown).

**S2.2.**
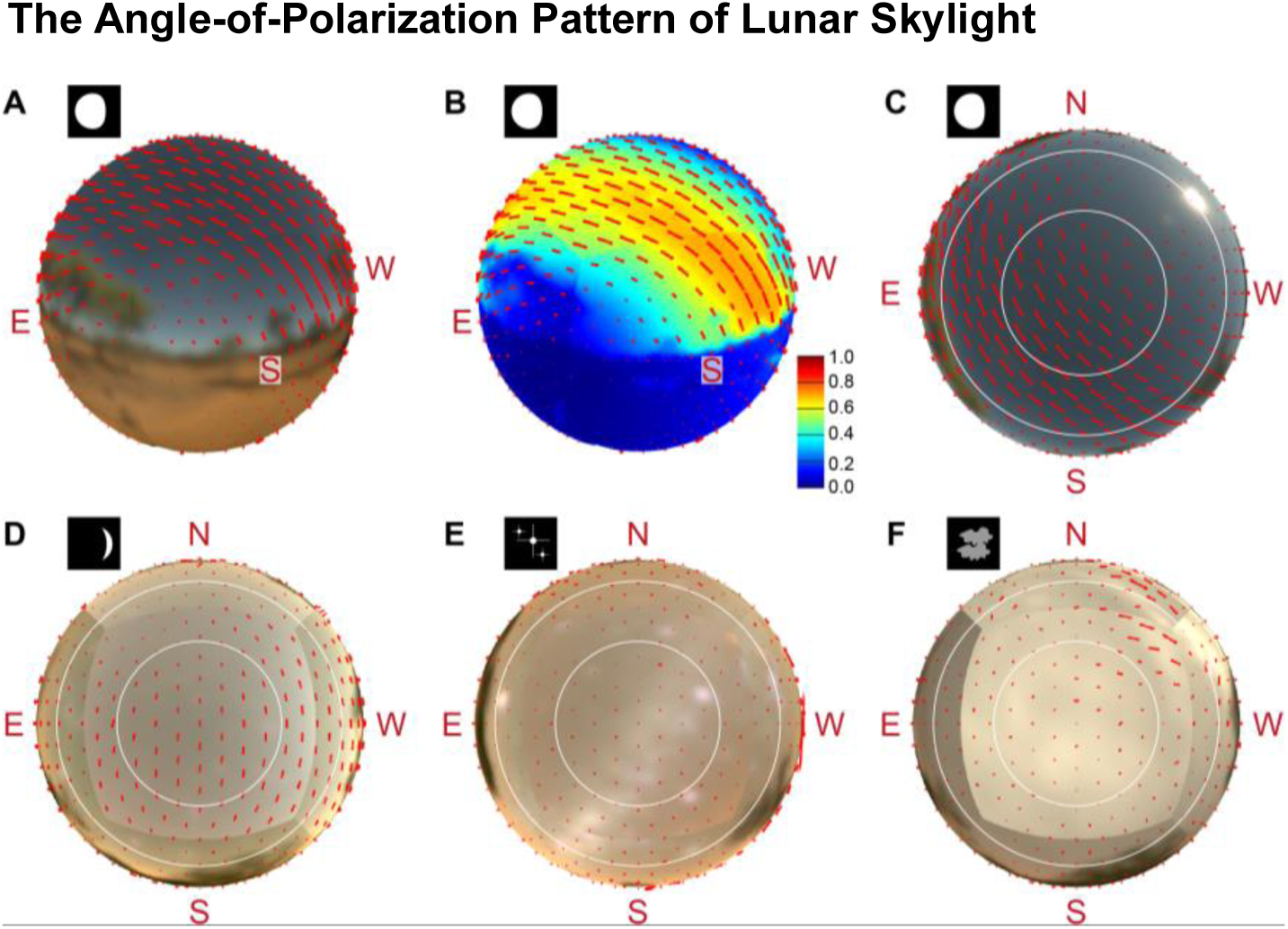
Estimated angles-of-polarization across the field of view measured. (A) shows an example image set for a waxing gibbous moon, projected onto a sphere of viewing angles. This sphere has been split into 642 equally-sized solid angles, and the angle of polarization calculated for each. Note that, because polarization was measured using a flat polarizing filter in before the fisheye lens in the light path, regions at the edge of two image sets, *e.g.* the zenith-facing and south-facing images, may appear to differ near to the boundary. (B) The length of these angle-of-polarization vectors has been scaled by their degree of polarization, which is less strongly affected by boundary discrepancies. (C) As for the degree of polarization figures, the angle-of-polarization pattern in the celestial hemisphere may be shown on an azimuth-elevation grid, although this somewhat distorts the true angle of polarization, which appears relative to the isolines of azimuth (vertical) and elevation (horizontal) rather than relative to the figure’s vertical and horizontal axes. (D) For the crescent moon, angles of polarization in adjacent regions were well aligned, although degree of polarization was far lower than for the other lunar phases. (E–F) In the absence of lunar skylight, angles of polarization in adjacent regions were not well aligned, suggesting that the larger degrees of polarization estimated in some regions are artefactual, or at least are not induced by scattering within the upper-atmosphere—as is the case for solar and lunar skylight polarization.

## References

Barta, A., Farkas, A., Száz, D., Egri, Á., Barta, P., Kovács, J., Csák, B., Jankovics, I., Szabó, G. and Horváth, G. (2014). Polarization transition between sunlit and moonlit skies with possible implications for animal orientation and Viking navigation: anomalous celestial twilight polarization at partial moon. Appl. Opt. 53, 5193–5204.

Batschelet, E. (1981). Circular statistics in biology. New York, NY, USA: Academic Press.

Bech, M., Homberg, U. and Pfeiffer, K. (2014). Receptive Fields of Locust Brain Neurons Are Matched to Polarization Patterns of the Sky. Curr. Biol. 24, 1–6.

Benn, C. R. and Ellison, S. L. (1998). Brightness of the night sky over La Palma. New Astron. Rev. 42, 503–507.

Berry, M. V., Dennis, M. R. and Lee, R. L. (2004). Polarization singularities in the clear sky. New J. Phys. 6, 1–14.

Boheman C. H. (1860). Coleoptera sammlade af J.A. Wahlberg i sydvestra Afrika. öfvers. Vetenskapsakad. Förh. Stockholm.

Bürkner, P.-C. (2017). Advanced Bayesian Multilevel Modeling with the R Package brms. J. Stat. Softw. 80, 1–18.

Carpenter, B., Gelman, A., Hoffman, M. D., Lee, D., Goodrich, B., Betancourt, M., Brubaker, M., Guo, J., Li, P. and Riddell, A. (2017). *Stan*: A Probabilistic Programming Language. J. Stat. Softw. 76.

Chiou, T.-H., Kleinlogel, S., Cronin, T. W., Caldwell, R., Loeffler, B., Siddiqi, A., Goldizen, A. and Marshall, N. J. (2008). Circular polarization vision in a stomatopod crustacean. Curr. Biol. 18, 1–6.

Dacke, M., Nordström, P., Scholtz, C. H. and Warrant, E. J. (2002). A specialized dorsal rim area for polarized light detection in the compound eye of the scarab beetle *Pachysoma striatum*. J. Comp. Physiol. A 188, 211–6.

Dacke, M., Byrne, M. J., Baird, E., Scholtz, C. H. and Warrant, E. J. (2011). How dim is dim? Precision of the celestial compass in moonlight and sunlight. Philos. Trans. R. Soc. B Biol. Sci. 366, 697–702.

Dacke, M., Byrne, M. J., Scholtz, C. H. and Warrant, E. J. (2004). Lunar orientation in a beetle. Proc. R. Soc. B 271, 361–365.

Dacke, M., Jundi, B., Smolka, J., Byrne, M. and Baird, E. (2014). The role of the sun in the celestial compass of dung beetles. Phil. Trans. R Soc. B 369, 20130036

Dacke, M., Nilsson, D.-E., Warrant, E. J., Blest, A. D., Land, M. F. and O’Carroll, D. C. (1999). Built-in polarizers form part of a compass organ in spiders. Nature 401, 470–474.

Dacke, M., Nordström, P. and Scholtz, C. H. (2003). Twilight orientation to polarised light in the crepuscular dung beetle *Scarabaeus zambesianus*. J. Exp. Biol. 206, 1535–1543.

Dacke, M., Nilsson, D.-E., Scholtz, C. H., Byrne, M. and Warrant, E. J. (2003). Insect orientation to polarized moonlight. Nature 424, 33.

Egri, Á., Farkas, A., Kriska, G. and Horváth, G. (2016). Polarization sensitivity in Collembola: an experimental study of polarotaxis in the water-surface-inhabiting springtail, *Podura aquatica*. J. Exp. Biol. 219, 2567–2576.

el Jundi, B., Foster, J. J., Byrne, M. J., Baird, E. and Dacke, M. (2015a). Spectral information as an orientation cue in dung beetles. Biol. Lett. 11, 20150656.

el Jundi, B., Warrant, E. J., Byrne, M. J., Khaldy, L., Baird, E., Smolka, J. and Dacke, M. (2015b). Neural coding underlying the cue preference for celestial orientation. PNAS 112, 11395–11400.

el Jundi, B., Foster, J. J., Khaldy, L., Byrne, M. J., Dacke, M. and Baird, E. (2016). A Snapshot-Based Mechanism for Celestial Orientation. Curr. Biol. 26, 1456–1462.

el Jundi, B., Smolka, J., Baird, E., Byrne, M. J., and Dacke, M. (2014). Diurnal dung beetles use the intensity gradient and the polarization pattern of the sky for orientation. J Exp Biol 217, 2422–2429.

Falchi, F., Cinzano, P., Duriscoe, D., Kyba, C. C. M., Elvidge, C. D., Baugh, K., Portnov, B. A., Rybnikova, N. A. and Furgoni, R. (2016). The new world atlas of artificial night sky brightness. Sci. Adv. 2, e1600377.

Foster, J. J., El Jundi, B., Smolka, J., Khaldy, L., Nilsson, D.-E., Byrne, M. J. and Dacke, M. (2017). Stellar performance: mechanisms underlying Milky Way orientation in dung beetles. Phil. Trans. R. Soc. B 372, 20160079.

Foster, J. J., Smolka, J., Nilsson, D.-E. and Dacke, M. (2018). How animals follow the stars. Proceedings. Biol. Sci. 285, 20172322.

Foster, J. J., Temple, S. E., How, M. J., Daly, I. M., Sharkey, C. R., Wilby, D. and Roberts, N. W. (2018). Polarisation vision: overcoming challenges of working with a property of light we barely see. Sci. Nat. 105, 27.

Freas, C. A., Narendra, A., Lemesle, C. and Cheng, K. (2017). Polarized light use in the nocturnal bull ant, *Myrmecia midas*. R. Soc. Open Sci. 4, 170598.

Frisch, K. von (1967). The dance language and orientation of bees. Cambridge, MA, US: Harvard University Press.

Gál, J., Horváth, G., Barta, A. and Wehner, R. (2001). Polarization of the moonlit clear night sky measured by full-sky imaging polarimetry at full Moon: Comparison of the polarization of moonlit and sunlit skies. J. Geophys. Res. 106, 22647–22653.

Glantz, R. M. and Schroeter, J. P. (2006). Polarization contrast and motion detection. J. Comp. Physiol. A 192, 905–914.

Hegedüs, R., Åkesson, S. and Horváth, G. (2007). Polarization patterns of thick clouds: overcast skies have distribution of the angle of polarization similar to that of clear skies. J. Opt. Soc. Am. A 24, 2347.

Henze, M. J. and Labhart, T. (2007). Haze, clouds and limited sky visibility: polarotactic orientation of crickets under difficult stimulus conditions. J. Exp. Biol. 210, 3266–3276.

Horváth, G., Barta, A. and Hegedüs, R. (2014). Polarization of the Sky. In Polarized Light in Animal Vision (ed. Horváth, G.), pp. 367–406. Berlin, Heidelberg: Springer Berlin Heidelberg.

Houpt, J. W. and Bittner, J. L. (2018). Analyzing thresholds and efficiency with hierarchical Bayesian logistic regression. Vision Res. 148, 49–58.

Johnsen, S., Kelber, A., Warrant, E., Sweeney, A. M., Widder, E. A., Lee, R. L. and Hernández-Andrés, J. (2006). Crepuscular and nocturnal illumination and its effects on color perception by the nocturnal hawkmoth Deilephila elpenor. J. Exp. Biol. 209, 789–800.

Kieffer, H. H. and Stone, T. C. (2005). The Spectral Irradiance of the Moon. Astron. J. 129, 2887–2901.

Kirwan, J. D. (2018). Spatial vision in diverse invertebrates. PhD thesis, Lund University, Lund, Sweden.

Knoblauch, K. and Maloney, L. T. (2012). Modeling Psychophysical Data in R. New York, NY: Springer New York.

Kyba, C. C. M., Ruhtz, T., Fischer, J. and Hölker, F. (2011). Lunar skylight polarization signal polluted by urban lighting. J. Geophys. Res. 116, D24106.

Labhart, T. (1996). How polarization-sensitive interneurones of crickets perform at low degrees of polarization. J. Exp. Biol. 199, 1467–1475.

Labhart, T. (1999). How polarization-sensitive interneurones of crickets see the polarization pattern of the sky: a field study with an opto-electronic model neurone. J. Exp. Biol. 202 (Pt 7), 757–70.

Labhart, T., Hodel, B. and Valenzuela, I. (1984). The physiology of the cricket’s compound eye with particular reference to the anatomically specialized dorsal rim area. J. Comp. Physiol. A 155, 289–296.

Menzel, R. and Snyder, A. W. (1974). Polarised light detection in the bee, Apis mellifera. J. Comp. Physiol. 88, 247–270.

Palmer, G. and Johnsen, S. (2015). Downwelling spectral irradiance during evening twilight as a function of the lunar phase. Appl. Opt. 54, B85.

Papi, F. and Pardi, L. (1963). On the Lunar Orientation of Sandhoppers (*Amphipoda Talitridae*). Biol. Bull. 124, 97–105.

Pfeiffer, K. and Homberg, U. (2007). Coding of Azimuthal Directions via Time-Compensated Combination of Celestial Compass Cues. Curr Biol 17, 960–965.

Pfeiffer, K., Negrello, M. and Homberg, U. (2011). Conditional perception under stimulus ambiguity: polarization- and azimuth-sensitive neurons in the locust brain are inhibited by low degrees of polarization. J. Neurophysiol. 105, 28–35.

Rossel, S. and Wehner, R. (1984). How bees analyse the polarization patterns in the sky. J. Comp. Physiol. A 154, 607–615.

Schmeling, F., Wakakuwa, M., Tegtmeier, J., Kinoshita, M., Bockhorst, T., Arikawa, K. and Homberg, U. (2014). Opsin expression, physiological characterization and identification of photoreceptor cells in the dorsal rim area and main retina of the desert locust, *Schistocerca gregaria*. J. Exp. Biol. 217, 3557–3568.

Schmidt-Koenig, K. (1990). The sun compass. Experientia 46, 336–342.

Shurcliff, W. A. (1955). Haidinger’s Brushes and Circularly Polarized Light. JOSA 45, 399.

Smolka, J., Baird, E., el Jundi, B., Reber, T., Byrne, M. J. and Dacke, M. (2016). Night sky orientation with diurnal and nocturnal eyes: dim-light adaptations are critical when the moon is out of sight. Anim. Behav. 111, 127–146.

Stavenga, D. G. (2010). On visual pigment templates and the spectral shape of invertebrate rhodopsins and metarhodopsins. J. Comp. Physiol. A Neuroethol. Sensory, Neural, Behav. Physiol. 196, 869–878.

Strutt, J. W. (1871). On the Light from the Sky, its Polarization and Colour. London, Edinburgh Dublin Philos. Mag. J. Sci. xxxvii, 107–120.

Tang, J., Zhang, N., Li, D., Wang, F., Zhang, B., Wang, C., Shen, C., Ren, J., Xue, C. and Liu, J. (2016). Novel robust skylight compass method based on full-sky polarization imaging under harsh conditions. Opt. Express 24, 15834.

Temple, S. E., Mcgregor, J. E., Miles, C., Graham, L., Miller, J., Buck, J., Scott-Samuel, N. E. and Roberts, N. W. (2015). Perceiving polarization with the naked eye: characterization of human polarization sensitivity. Proc. R. Soc. B 282,.

Templin, R. M., How, M. J., Roberts, N. W., Chiou, T.-H. and Marshall, J. (2017). Circularly polarized light detection in stomatopod crustaceans: a comparison of photoreceptors and possible function in six species. J. Exp. Biol. 220, 3222–3230.

Tibbs, A. B., Daly, I. M., Bull, D. R. and Roberts, N. W. (2018). Noise creates polarization artefacts. Bioinspir. Biomim. 13, 015005

Wang, X., Gao, J., Fan, Z. and Roberts, N. W. (2016). An analytical model for the celestial distribution of polarized light, accounting for polarization singularities, wavelength and atmospheric turbidity. J. Opt. 18, 065601.

Warrant, E. J. (1999). Seeing better at night: Life style, eye design and the optimum strategy of spatial and temporal summation. Vis. Res. 39, 1611–1630.

Wehner, R. (2001). Polarization vision—a uniform sensory capacity? J. Exp. Biol. 204, 2589–2596.

Zeil, J., Ribi, W. A. and Narendra, A. (2014). Polarisation vision in ants, bees and wasps. In Polarized Light in Animal Vision (ed. Horváth, G.), pp. 41–60. Berlin, Heidelberg: Springer Berlin Heidelberg.

## Supplementary References

Dacke, M., Nordström, P., Scholtz, C. H. and Warrant, E. J. (2002). A specialized dorsal rim area for polarized light detection in the compound eye of the scarab beetle Pachysoma striatum. J. Comp. Physiol. A 188, 211–6.

Dacke, M., Nordström, P. and Scholtz, C. H. (2003). Twilight orientation to polarised light in the crepuscular dung beetle Scarabaeus zambesianus. J. Exp. Biol. 206, 1535–1543.

Lord, N. P., Plimpton, R. L., Sharkey, C. R., Suvorov, A., Lelito, J. P., Willardson, B. M. and Bybee, S. M. (2016). A cure for the blues: Opsin duplication and subfunctionalization for short-wavelength sensitivity in jewel beetles (Coleoptera: Buprestidae). BMC Evol. Biol. 16, 107.

