## Supplementary figures and images for "Orienting to Polarized Light at Night—Matching Lunar Skylight to Performance in a Nocturnal Beetle"

### Supplementary file 1

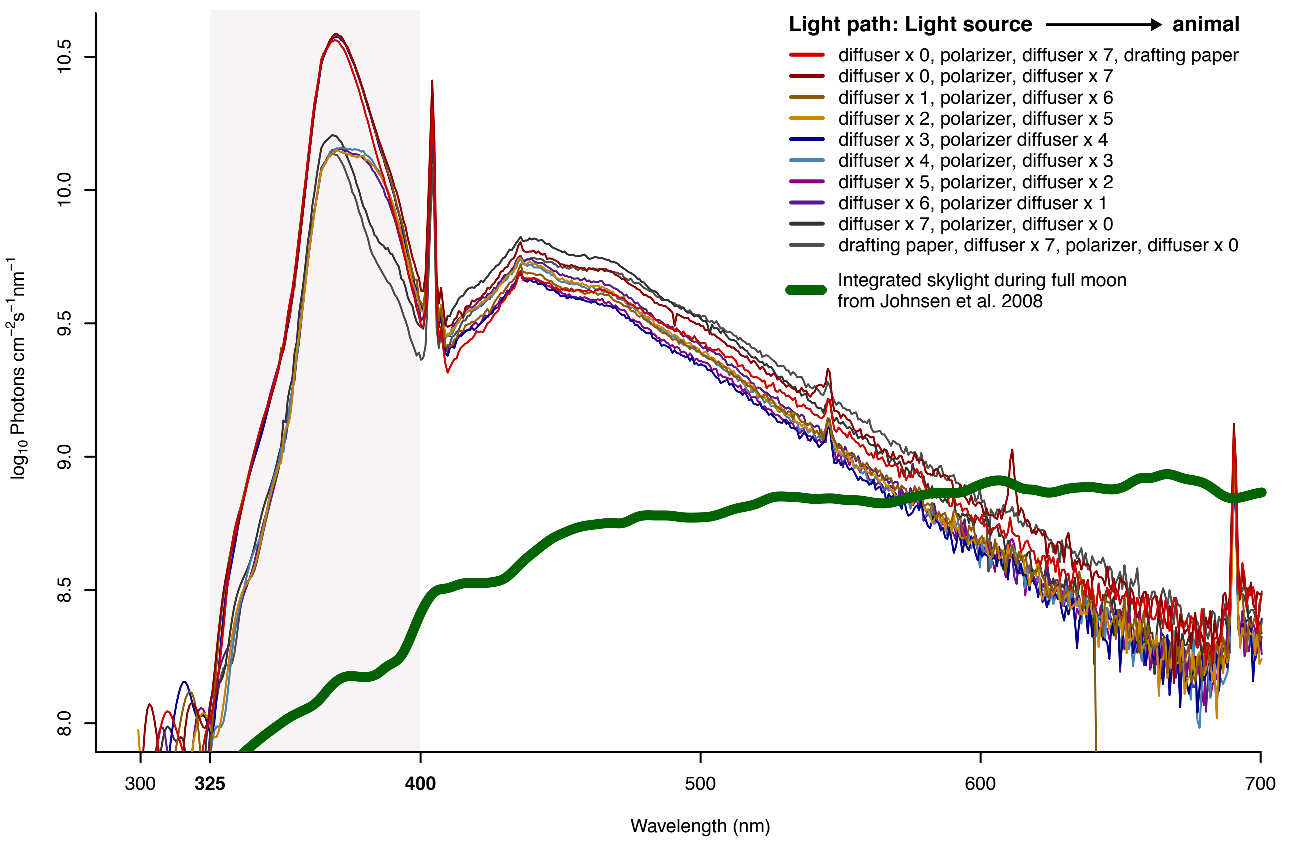

### Supplementary file 2

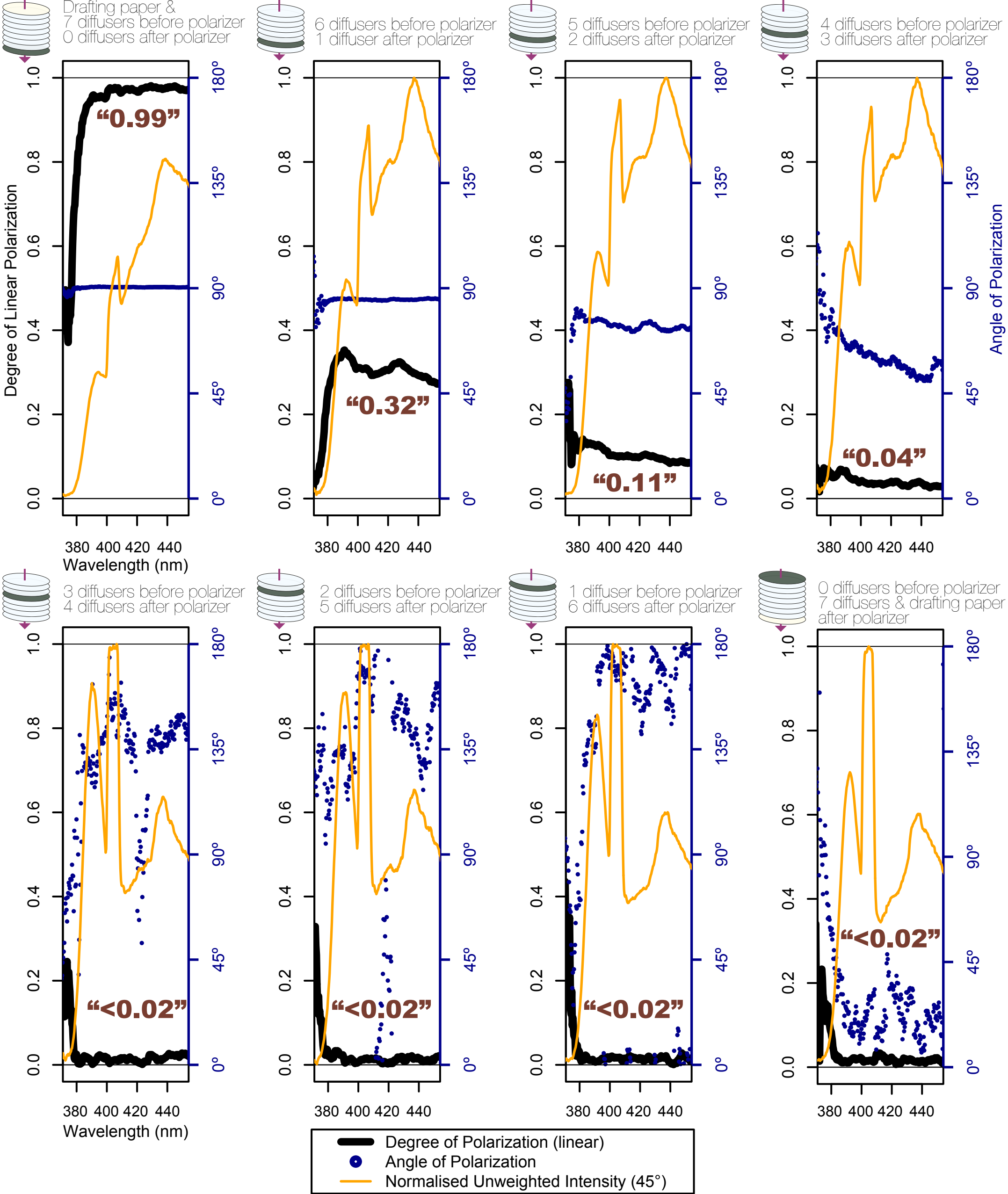

### Supplementary file 3

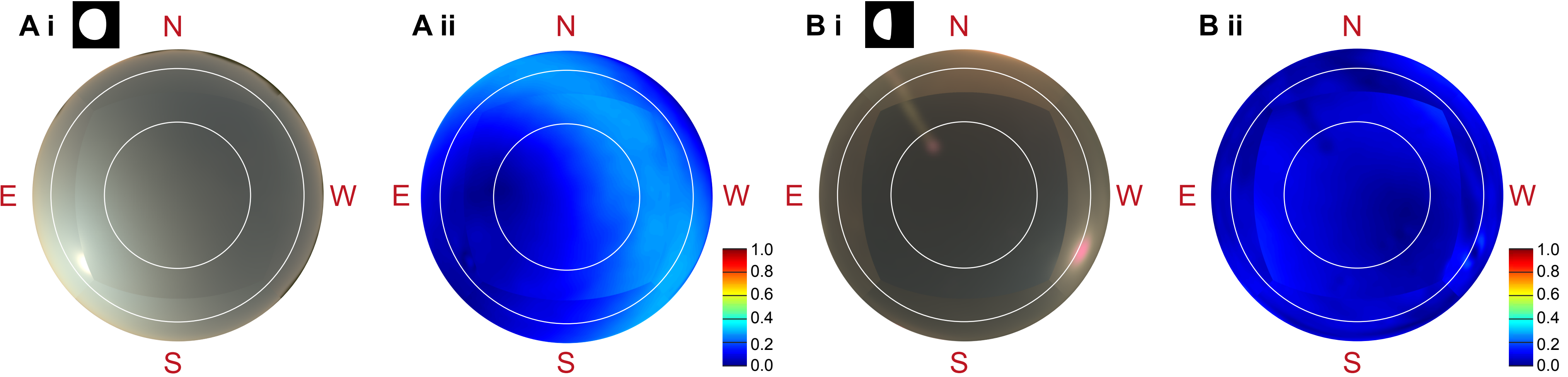

### Supplementary file 4

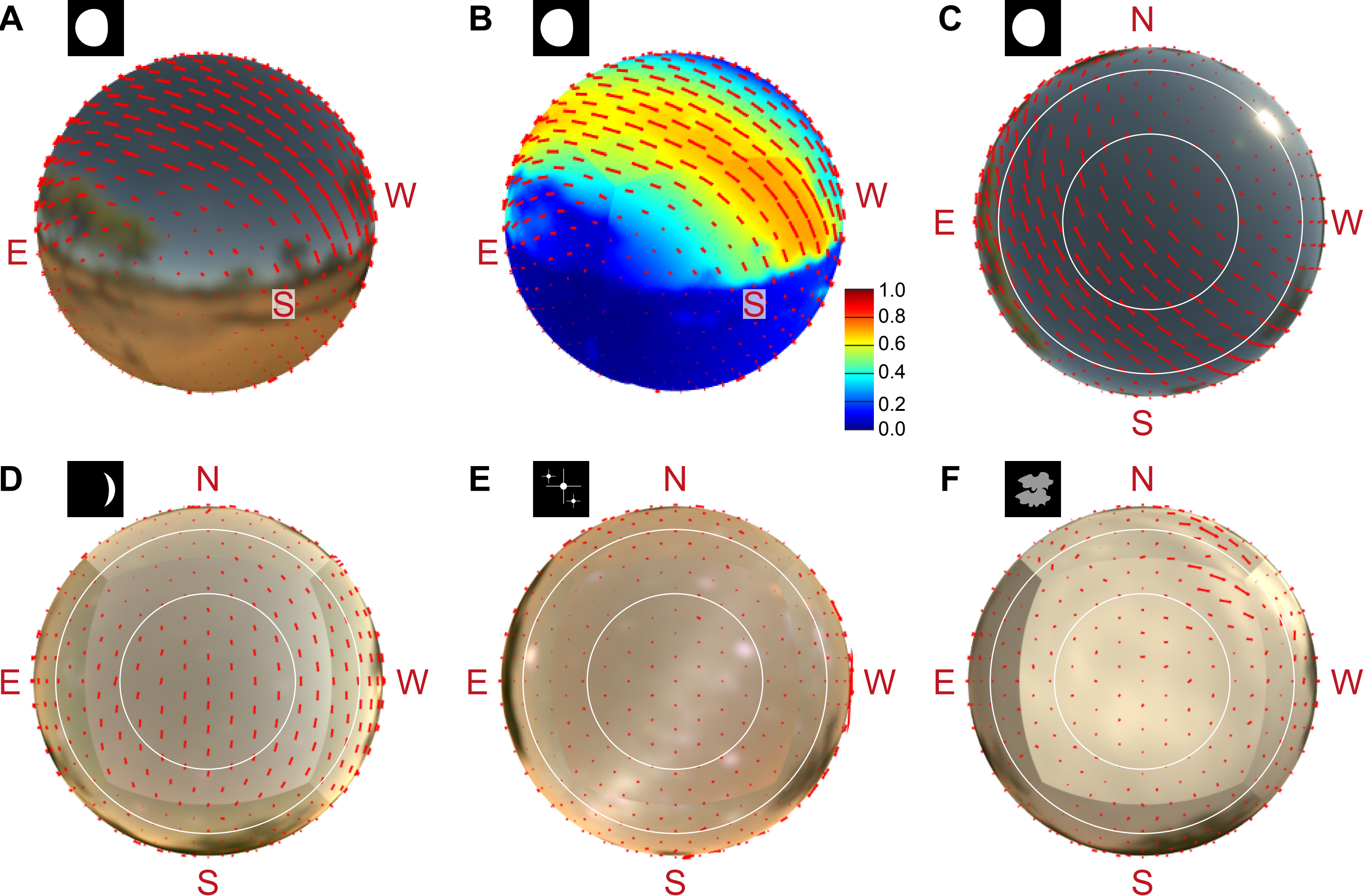
